# Studies of Bending Effects of Microvilli of Leukocyte on Rolling Adhesion

**DOI:** 10.1101/322198

**Authors:** Tai-Hsien Wu, Dewei Qi

**Author notes:** Correspondence. Address reprint requests to Dewei Qi, 1903 W. Michigan Ave., Kalamazoo, MI 49009.

## Abstract

It has been widely acknowledged that further understanding about the cell adhesion (e.g., leukocyte rolling adhesion) can help us gain more knowledge about the causes of relevant diseases and design more effective treatments and diagnoses. Although recent simulation studies considered the deformability of the leukocytes, most of them, however, did not consider the bending deformation of microvilli. In this paper, an advanced leukocyte model based on an immersed boundary lattice-Boltzmann lattice-spring model (LLM) and an adhesive dynamics (AD) is presented in details. The flexural stiffness of microvilli is introduced into the model for simulations of leukocyte rolling adhesion. This innovative model is applied to investigate the influences of bending deformation of microvilli on the process of leukocyte rolling adhesion and the underlying mechanism at different shear rates. It is demonstrated that the bending deformation of microvilli can be influenced by the flexural stiffness of microvilli and shear rates, resulting in the different rolling velocity of leukocytes, number of receptor-ligand bonds, and bond forces. The findings clearly indicate that the bending of microvilli plays a crucial role in the dynamics of leukocyte adhesion.

## INTRODUCTION

Cell adhesion plays important roles in various in-vivo and in-vitro binding applications. For example, chimeric antigen receptor (CAR) T-cells rely on binding to cancer cells, under various shear rates, in order to slow or halt potential cancer growth (1, 2). A more general example is leukocytes rolling along the inflammation-activated vascular endothelium in blood flow, which is a part of the human immune response. This rolling process is mediated by selectin-ligand bonds, such as P-selectin–P-selectin glycoprotein ligand-1 (PSGL-1) bonds. Many experimental and numerical studies have confirmed that the dynamics of adhesion are mainly mediated by the physical chemistry of adhesion molecules, such as the association and dissociation rates of adhesion bonds (3, 4), selectin/ligand density, and the local circulatory environment including flow shear rate (3, 5).

In the earliest attempts of simulation, leukocytes were modeled by rigid spheres. However, cellular material properties such as cell deformation play a crucial role in the dynamics of cell adhesion. Khismatullin and Truskey (6, 7) first used a compound viscoelastic drop to model neutrophils and monocytes to investigate the effects of cell deformability in a parallel-plate flow chamber. Almost at the same time, Jadhav et al. (8) used a neo-Hookean membrane as a leukocyte, coupled with the immersed boundary method (IBM), to investigate the influence of cell deformability on leukocyte rolling adhesion. Their results indicated that cells with deformations roll much slower and are relatively more stable than those without deformation in shear flow, facts consistent with the experimental findings (9, 10). Since then, numerous simulation studies continue to address the process of cell deformation during cellular adhesion (11-16).

Another critical cellular property is microvillus deformation. Microvilli are fingerlike projections on the surface of leukocytes. Since most adhesion molecules (e.g., PSGL-1) are located at the tips of microvilli, microvillus deformation also affects cellular adhesion. Hence, several simulation studies were performed to examine the influence of the extensional deformation of microvilli on the dynamics of cell adhesion. Khismatullin and Truskey (6) combined the microvillus spring with the receptor-ligand bond spring to express the total extensional deformation of microvilli and adhesive bonds. Later, Caputo and Hammer (17) improved Hammer and Apte’s pioneering adhesive dynamics (AD) (3) by using an elastic spring and a viscous model to approximate the microvilli elastic and viscous responses to a small force and a large force, respectively. This idea was first proposed by Shao et al. (18). Pawar et al. (19) further improved Caputo and Hammer’s model by replacing the hard spherical body with a deformable neo-Hookean membrane. Recently, Pospieszalska et al. combined the existing model for microvillus tether formation with a Kelvin-Voigt viscoelastic model for microvillus extension (20, 21). However, these studies were limited to microvillus extensional deformation.

In fact, when a force tangential to the cell surface is exerted on a leukocyte microvillus, it will be bent. This phenomenon was observed in experiments (22). Similar to the extensional deformation of microvilli, the bending of microvilli due to their flexibility can potentially influence the binding between selectins and ligands. Munn et al. (23) suggested that when microvilli are flattened along the cell surface, the receptors at the bases of microvilli (instead of at the tips of microvilli) may obtain additional adhesion with ligands on the endothelial cells or on the ligand-coated substrate. While Pospieszalska et al. first simulated the pivoting of microvilli around their bases, they did not address the flexural stiffness of microvilli (20, 21). Based on the importance of flexibility in microvilli, Yao and Shao (24, 25) experimentally measured the flexural stiffness of the microvilli on lymphocytes and neutrophils under small deformation of the microvilli. They reported that the values of the flexural stiffness of lymphocyte and neutrophil microvilli are 4*pN/μm* and 7*pN/μm*, respectively (24, 25). Simon et al. (26) conducted another experiment to measure the flexural stiffness of neutrophil microvilli under a large deformation (0.5 − 1*μm*) and found that the flexural stiffness is 5*pN/μm*.

To take account of the bending effects of microvilli, employing the measured data of microvillus flexural stiffness identified by Yao and Shao (24, 25), Wu and Qi (27) recently used the immersed boundary lattice-Boltzmann lattice-spring model (LLM) combined with adhesive dynamics simulations (3) to study the roles of flexural stiffness on cell rolling and adhesion at a shear rate of *γ* = 50*s*^−1^. The results showed that the flexural stiffness has a profound effect on rolling velocity and adhesion bonding force. In their studies, they decomposed each individual adhesion bonding force into two local coordinate-based components: parallel (for extension) and perpendicular (for bending). Their findings revealed that the increasing total local coordinate-based adhesion bonding force is mainly due to an increase in the perpendicular component. The flexibility of microvilli aids not only in expediting contact frequency with the substrate, but also in flattening of microvilli once the contact is established. As a result, the flexibility facilitates the formation of adhesive bonds, confirming Munn et al’s speculation (23).

This paper advances the study of microvillus flexibility in leukocyte rolling adhesion under different shear rates. In simulations, the statistical ensemble average results of individual microvilli are computed and used to understand the dynamic adhesive behavior, which is difficult to observe in experiments because of limited spatial and temporal resolution currently available. The results suggest that the flexural stiffness of microvilli plays a key role in altering the rolling velocity of leukocytes at different shear rates. As the flexural stiffness of microvilli increases or its flexibility decreases, the leukocytes roll faster or detach from the selectin-coated substrate more easily. This phenomenon is apparent even at a higher shear rate.

The same simulation method developed by Wu and Qi (27) is adopted in the present work. This method is comprised of five numerical models: (1) lattice Boltzmann method (LBM) for the Navier-Stokes flow (28-37); (2) coarse-grained cell model (CGCM) for the viscoelastic leukocyte membrane (13, 38-41); (3) lattice spring model (LSM) for the flexible microvilli (42); (4) immersed boundary method (IBM) for the fluid-cell interactions (43); (5) adhesive dynamics (AD) for stochastic binding between P-selectin and PSGL-1 (3). The method with each model is presented in details in the next section, and the contents were neither reported nor published before.

## COMPUTATION METHOD

A lattice-Boltzmann lattice-spring method (LLM) (27, 42, 44-46) combined with the AD is employed. In this method, the LBM is used as a fluid solver to simulate the Navier-Stoke flows, the CGCM (38) is utilized to simulate motion of leukocytes, and the LSM (27) is exploited to mimic the deformation of microvilli while the fluid-solid interactions are treated by IBM. Further, the AD is employed to obtain the adhesive bonding forces between leukocytes and selectin-coated substrates.

### Lattice Boltzmann method

The motion of Newtonian fluid is governed by the continuity and Navier-Stokes equations as

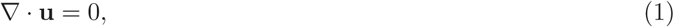

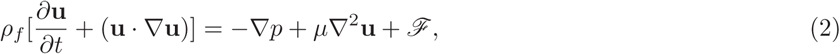

where **u** is the fluid velocity; *ρ_f_* the fluid density; *p* the pressure; *μ* the fluid dynamic viscosity; and ℱ the force source term. In this study, the partial differential equations (Eqs. 1 and 2) are not solved directly. Instead, the LBM is used to obtain fluid flow behavior. Previous work has demonstrated that the solution of the LBM is equivalent to that of the Navier-Stokes equations when the Mach number is smaller than 0.3 (30). Importantly, the kinetic nature of the LBM makes it more suitable for simulating the multi-component flows such as blood (47-51). More features of the LBM can be referred to previous studies (35).

The multiple-relaxation-time (MRT) lattice Boltzmann equation (52) in a D3Q19 lattice model with a forcing term is used and expressed as

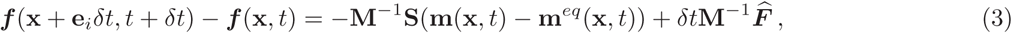

where ***f***(**x**, *t*) and **m**(**x**, *t*) are the fluid distribution functions at position **x** and time *t* in the velocity and moment spaces, respectively; m^eq^ denotes the equilibrium distribution function in moment space; *δt* represents the time interval; **e***_i_* is the discrete velocity set where *i* ∈ {0,1, 2,..., 18} are the discrete directions; **S** is the diagonal collision matrix; **M** is the transformation matrix (given in Ref. (52) Appendix) which transfers the distribution functions from the velocity space into the moment space; and 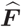 is the moment of the forcing term in the moment space.

In the D3Q19 model, the discrete velocity set **e***_i_* is written as:

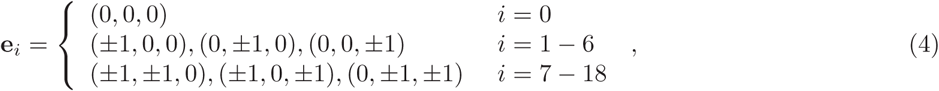

and diagonal collision matrix **S** is written as:

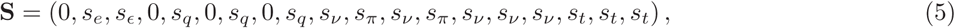

while shear and bulk viscosities are given by

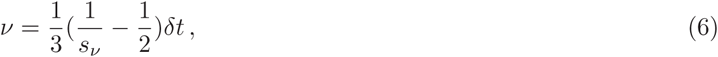

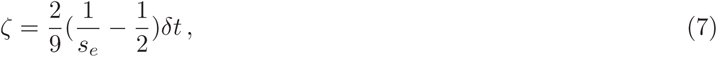

where *s_v_* is related to the shear viscosity. As suggested by Pan et al. (53), the other relaxation rates are set as

*s_e_* = *s_ε_ = s_π_ =s_v_* and 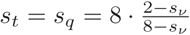.

The distribution functions in the moment space m and the corresponding equilibria **m***^eq^* are written by

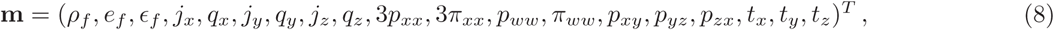

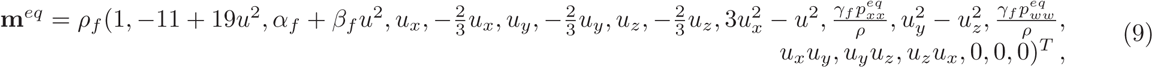

where *e_f_* and *e_f_* represent energy and energy squared; *j_x_,_y_,_z_* are components of the momentum; *q_x_,_y_,_z_* are components of the heat flux; *p_xy_,_yz_,_zx_* are the symmetric and traceless strain-rate tensor; *n_xx_,_ww_* are the fourth order moments and *t_x_,_y_,_z_* are the third order moments (52); 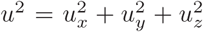 denotes the fluid velocity squared; and the three parameters *α_f_* = 3, 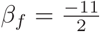, and 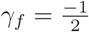 are chosen.

Furthermore, according to the work by Guo et al. (54, 55), the moment of the forcing term in the moment space 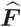 can be written as,

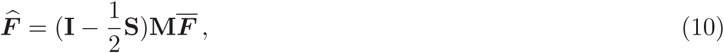

where **I** is the unity matrix; 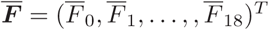 and

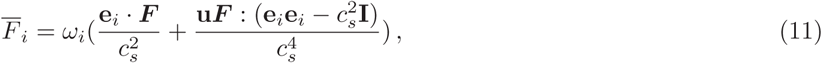

where ***F*** is the body force; 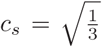 is the sound speed; *ω_i_* is the weight associated with the lattice model and defined by

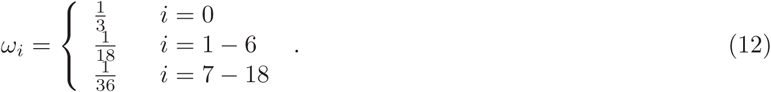

The fluid velocity **u** is given by

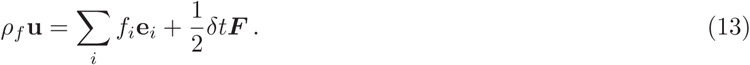

All parameters used in the LBM are given in Table 1 and Table 2.

**Table 1:**
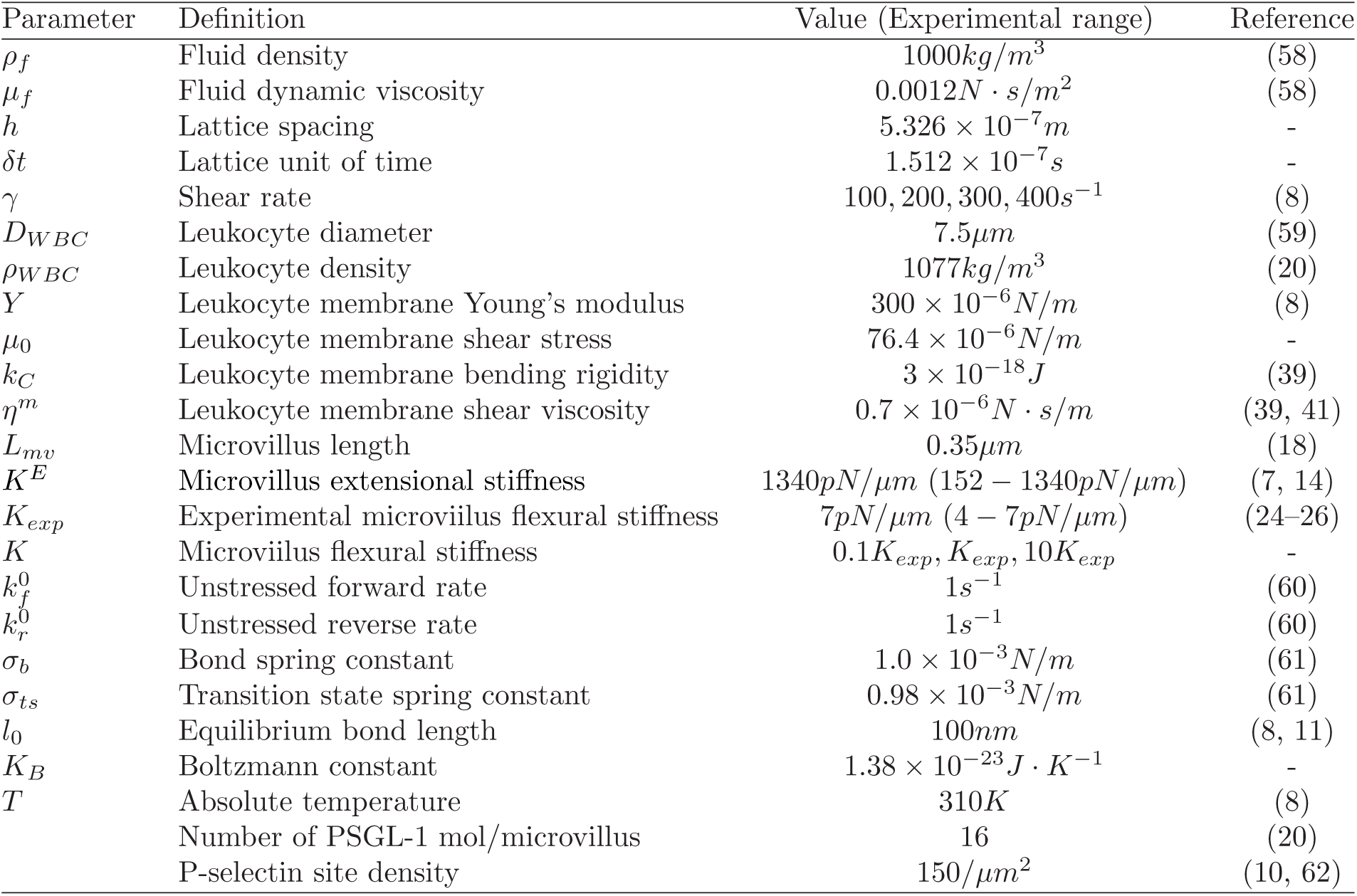
The simulation dimensional parameters used in this study.

| Parameter | Definition | Value (Experimental range) | Reference |
| --- | --- | --- | --- |
| $\rho_f$ | Fluid density | $1000kg/m^3$ | (58) |
| $\mu_f$ | Fluid dynamic viscosity | $0.0012N \cdot s/m^2$ | (58) |
| $h$ | Lattice spacing | $5.326 \times 10^{-7}m$ | - |
| $\delta t$ | Lattice unit of time | $1.512 \times 10^{-7}s$ | - |
| $\gamma$ | Shear rate | $100, 200, 300, 400s^{-1}$ | (8) |
| $D_{WBC}$ | Leukocyte diameter | $7.5\mu m$ | (59) |
| $\rho_{WBC}$ | Leukocyte density | $1077kg/m^3$ | (20) |
| $Y$ | Leukocyte membrane Young's modulus | $300 \times 10^{-6}N/m$ | (8) |
| $\mu_0$ | Leukocyte membrane shear stress | $76.4 \times 10^{-6}N/m$ | - |
| $k_C$ | Leukocyte membrane bending rigidity | $3 \times 10^{-18}J$ | (39) |
| $\eta^m$ | Leukocyte membrane shear viscosity | $0.7 \times 10^{-6}N \cdot s/m$ | (39, 41) |
| $L_{mv}$ | Microvillus length | $0.35\mu m$ | (18) |
| $K^E$ | Microvillus extensional stiffness | $1340pN/\mu m$ ($152 - 1340pN/\mu m$) | (7, 14) |
| $K_{exp}$ | Experimental microvillus flexural stiffness | $7pN/\mu m$ ($4 - 7pN/\mu m$) | (24–26) |
| $K$ | Microvillus flexural stiffness | $0.1K_{exp}, K_{exp}, 10K_{exp}$ | - |
| $k_f^0$ | Unstressed forward rate | $1s^{-1}$ | (60) |
| $k_r^0$ | Unstressed reverse rate | $1s^{-1}$ | (60) |
| $\sigma_b$ | Bond spring constant | $1.0 \times 10^{-3}N/m$ | (61) |
| $\sigma_{ts}$ | Transition state spring constant | $0.98 \times 10^{-3}N/m$ | (61) |
| $l_0$ | Equilibrium bond length | $100nm$ | (8, 11) |
| $K_B$ | Boltzmann constant | $1.38 \times 10^{-23}J \cdot K^{-1}$ | - |
| $T$ | Absolute temperature | $310K$ | (8) |
|  | Number of PSGL-1 mol/microvillus | 16 | (20) |
| | P-selectin site density | $150/\mu m^2$ | (10, 62) |

**Table 2:** The simulation dimensionless parameters used in this study.

| Parameter | Definition | Value |
| --- | --- | --- |
| **LBM** |  |  |
| $\rho_f$ | Fluid density | 1 |
| $\nu_f$ | Fluid kinematic viscosity | 0.64 |
| $\gamma$ | Shear rate | $(1.512, 3.024, 4.536, 6.048) \times 10^{-5}$ |
| $(N_x, N_y, N_z)$ | Simulation box | (32, 48, 32) |
| $(s_e, s_\epsilon, s_\pi, s_\nu, s_t, s_q)$ | Relaxation rates | (0.41, 0.41, 0.41, 0.41, 1.67, 1.67) |
| $(\alpha_f, \beta_f, \gamma_f)$ | Three free parameters in the MRT | (3, -5.5, -0.5) |
| **CGCM** |  |  |
| $N_p$ | Number of solid nodes | 714 |
| $\rho_{WBC}$ | Leukocyte density | 1.077 |
| $Y$ | Leukocyte membrane Young’s modulus | 0.0454 |
| $\mu_0$ | Leukocyte membrane shear stress | 0.0116 |
| $k_C$ | Leukocyte membrane bending rigidity | 0.0016 |
| $(\eta^T, \eta^C)$ | dissipative parameters | $(1.9896 \times 10^{-4}, 6.6321 \times 10^{-5})$ |
| $m$ | Exponent in POW | 6 |
| $k_F$ | FENE coefficient | $2.298 \times 10^{-3}$ |
| $k_A$ | Global area constant | 0.294 |
| $k_D$ | Local area constant | 0.00589 |
| $k_V$ | Volume constant | 0.294 |
| $k_B$ | Bending coefficient | 0.00185 |
| $R^C$ | Cut-off radius of the repulsive potential | 0.055 |
| $\sigma_R$ | Well depth of the repulsive potential | 0.05 |
| $\epsilon_R$ | Distance where repulsive potential equals zero | $2 \times 10^{-5}$ |
| **LSM** |  |  |
| $N_{mv}$ | Number of microvilli | 714 |
| $L_{mv}$ | Microvillus length | 0.6571 |
| $k_s^{mv}$ | Microvillus spring constant | 0.203 |
| $k_a^{mv}$ | Microvillus angle constant | $(4.58 \times 10^{-5}, 4.58 \times 10^{-4}, 4.58 \times 10^{-3})$ |
| **AD** |  |  |
| $k_f^0$ | Unstressed forward rate | $1.51287 \times 10^{-7}$ |
| $k_r^0$ | Unstressed reverse rate | $1.51287 \times 10^{-7}$ |
| $\sigma_b$ | Bond spring constant | 0.151495 |
| $\sigma_{ts}$ | Transition state spring constant | 0.14998 |
| $l_0$ | Equilibrium bond length | 0.1877 |
| $K_B T$ | Thermal energy | $2.2858 \times 10^{-6}$ |
|  | P-selectin site density | 43 |
| $\Delta t$ | Time interval for adhesion determination | 1000 |
| $A_L n_L$ | Number of PSGL-1 accessible to each receptor | 5 |

### Immersed boundary method

In the LBM, the fluid particles are in regular Eulerian grids, whereas the solid particles are in Lagrangian grids. Therefore, a solid grid may not coincide with its adjacent fluid grids. The IBM is used to achieve non-slip boundary condition in the fluid-solid interfaces. For the purpose, a discrete Dirac delta function *δ^D^* is taken to interpolate the fluid velocity at the position of a solid boundary grid from its surrounding fluid grids. The discrete Dirac delta function (43) is written by

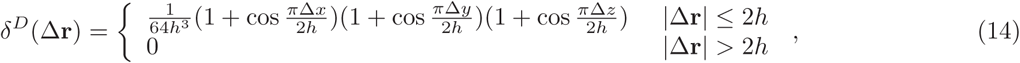

where *h* is the fluid grid spacing and ∆**r** = (∆*x*, ∆*y*, ∆*z*) is the distance between the positions of the solid boundary grid and its surrounding fluid grids.

To present the IBM, the solid boundary domain Γ and fluid boundary domain Π are defined. The solid boundary domain Γ is constituted by all the solid grids on the surface, and the fluid boundary domain Π is defined as a spherical volume of a radius of 2*h* centered at a solid boundary grid position **r***^b^*. The un-forced fluid velocity **u**^*^ at **r***^b^* is represented by

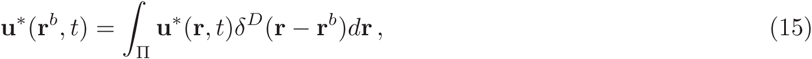

where **r** is a variable and goes all the positions of the fluid grids in the fluid domain Π during the integration. Due to the no-slip boundary condition, the forced fluid velocity **u**(**r***^b^*, *t*) should be equal to the solid velocity **U***^b^*(**r***^b^*, *t*), as follows:

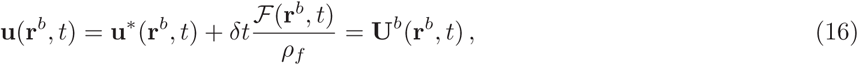

where 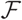 is the fluid-solid interaction force exerted on the fluid. Therefore, the interaction force 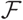 exerted on the fluid at the solid boundary position can be expressed by

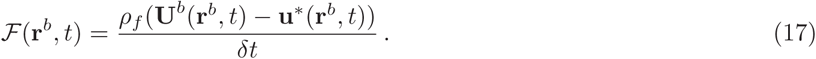

Thus, the interaction force acting on the solid grids by the surrounding fluid is given by

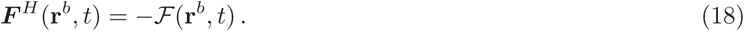

The discrete Dirac delta function is utilized again to distribute the interaction force 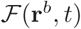 to the surrounding fluid grids by

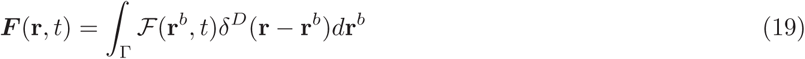

where ***F***(**r**, *t*) is the distributed body force and can be used in Eq. 11.

### Coarse-grained cell model

A leukocyte can be simulated by a coarse-grained cell model (CGCM), presented by Fedosov et al. (38), in which the leukocyte is assumed to be a spherical membrane. The membrane is considered as a two-dimensional triangulated network. An open source MATLAB mesh generator presented by Persson et al. (56) was used in this study for creating meshes.

In the triangulated network, *N_p_* denotes the number of solid grids, *N_b_* the number of edges, and *N_e_* the number of triangles. Four types of potential energies are calculated between solid grids. The total potential energy of the membrane can be written as

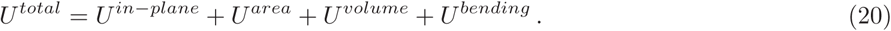

The in-plane potential energy term *U^in–plane^* has several formulas constituted by two-body and three-body energies. In this study, an edge is considered as a non-linear spring, whose energy is constituted by the combination of the finitely extensible nonlinear elastic (FENE) and power law (POW) potential energies, as follows:

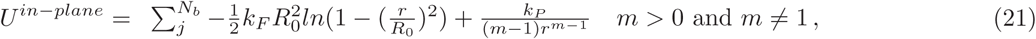

where *k_F_* is a constant coefficient of the FENE potential; *R*_0_ and *r* are the maximum and instant distances between two solid grids, respectively; *k_P_* is the coefficient of the POW potential energy; and, *m* is the exponent of the power law. Apparently, *U^in–plane^* is only a two-body energy of a non-linear spring.

The area conservation and volume conservation energies, *U^area^* and *U^volume^*, are the three-body energies which exist in a triangle constituted by three neighboring solid grids. The area and volume conservation energies are defined as follows:

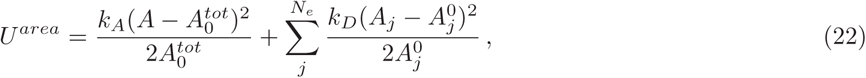

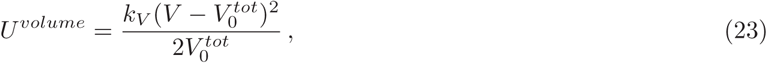

where *k_A_*, *k_D_*, and *k_V_* are the global area, local area, and volume constraint constants, respectively. The term *A* and *V* represent the instantaneous entire area and volume, respectively, while 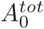 and 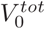 are the initial total area and volume; 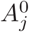 and *A_j_* are the initial and instantaneous local area of the *j*th triangle.

Furthermore, the bending energy *U^bending^* exists between two adjacent triangles (four adjacent solid grids) and is given by the following equation:

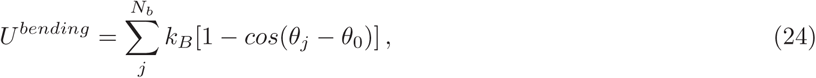

where *k_B_* is the bending coefficient; and *θ_j_* and *θ*_0_ are the instantaneous and initial angles, respectively, between two adjacent triagnels which have the common edge *j*.

The total elastic force 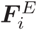 exerted on the *i*th grid in the CGCM is computed by

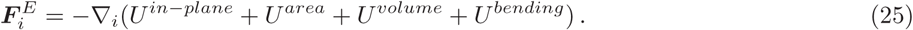

The gradient is calculated analytically, and the answers are used in the code.

The CGCM (13, 38-41) also addresses the membrane viscosity by adding the dissipative 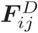 for each spring (edge) as

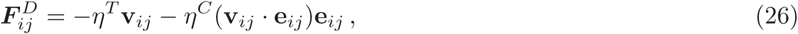

where *η^T^* and *η^C^* are dissipative coefficients, respectively; and **v***_ij_* and **e***_ij_* are the relative velocity and unit vector, respectively, between the *i*th and *j*th grids. The membrane shear viscosity *η^m^* is related to the dissipative parameters (*η^T^* and *η^C^*), as follows:

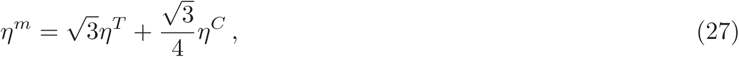

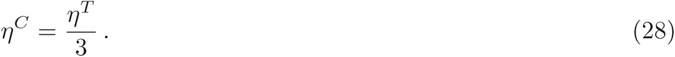

A repulsive force is added on leukocytes between the leukocytes and substrate in the Z-direction (perpendicular to the substrate) when the leukocytes are close to the substrate. The repulsive force is borrowed from the gradient of the Lennard-Jones potential (*U^LJ^*) with negative sign, as follows:

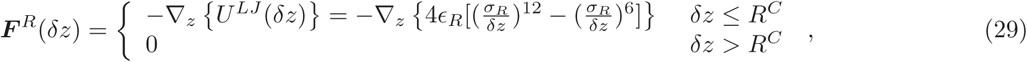

where *δz* is the distance above the substrate; *R^C^* is the cut-off radius; *ε_R_* is the well depth of the potential; and *σ_R_* is the distance between the substrate surface and the position where *U^LJ^* = 0. Therefore, the total force exerted on the *i*th solid grid is given by

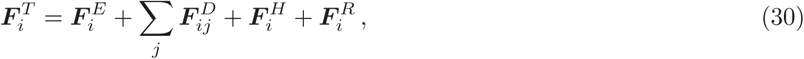

where ***F****^T^*, ***F****^E^*, ***F****^D^*, ***F****^s^*, and ***F****^R^* denote the total, elastic, dissipative, hydrodynamic, and repulsive forces, respectively; and *j* denotes the grid nearby the *i*th solid grid. Once the total force of a solid grid is obtained, the leap-frog algorithm is used to obtain the position and velocity of each solid grid during the simulation run.

Next, six parameters (*k_F_*, *k_P_*, *k_A_*, *k_D_*, *k_V_*, and *k_B_*) must be determined in the CGCM. First, the potential coefficient of POW *k_P_* can be described in terms of *k_F_* since the attractive term should be equal to the repulsive term at the equilibrium point in *U^in–plane^*. By setting the exponent of the power law *m* = 6 and *R*_0_ = 1.75*r*, the potential coefficient of power law *k_P_* can be written by

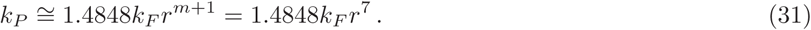

Secondly, according to Fedosov et al. (38, 39), the shear stress *μ*_0_ based on the *U^in–plane^* is given by

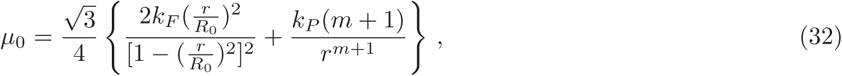

and the compression modulus *κ* of cell is given by

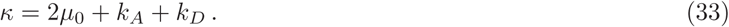

The linear Young’s modulus *Y* and Poisson’s ratio *v^p^* of the cell are expressed by

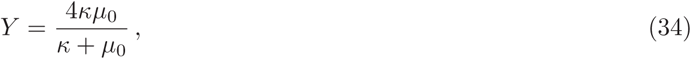

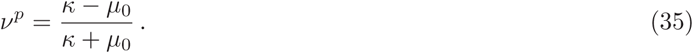

To achieve incompressibility and to make *k_A_* + *k_D_* >> *μ*_0_, Fedosov et al. showed that a nearly incompressible membrane is achieved when *k_A_* + *k_D_* = 500*μ*_0_. The assumption is followed. In addition, the bending coefficient *k_B_* in the CGCM is given by

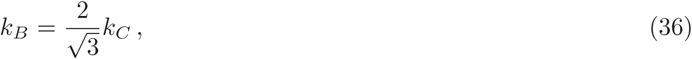

where *k_C_* is the bending rigidity. All parameters used in the CGCM are collected in Table 1 and Table 2.

### Lattice spring model

Wu and Qi (27) applied the LSM (42) to mimic the deformation of the leukocyte microvilli. As shown in Fig. 1, a microvillus is modeled by a spring (vector **r***_ji_*.) with its end fixed at the base grid *i* on the cell surface and with the other end *j* as the microvillus tip. Note that the base grids in the LSM are also the solid grids in the CGCM. There are two types of deformation of microvilli. First, the tip grid *j* of the spring can be extended or compressed with respect to the base grid *i*. The potential energy between grids *i* and *j* responsible for the spring extension or compression 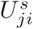 is given by

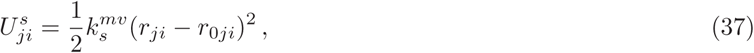

where *r_ji_* and *r*_0_*_ji_* are the instantaneous and initial microvillus lengths, respectively; and 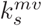 is the extensional spring constant. Note that each microvillus has the same initial length (i.e., *r*_0_*_ji_* = *L_mv_*, where *L_mv_* = 0.35*μm* is the average experimental value of microvillus length).

**Figure 1:**
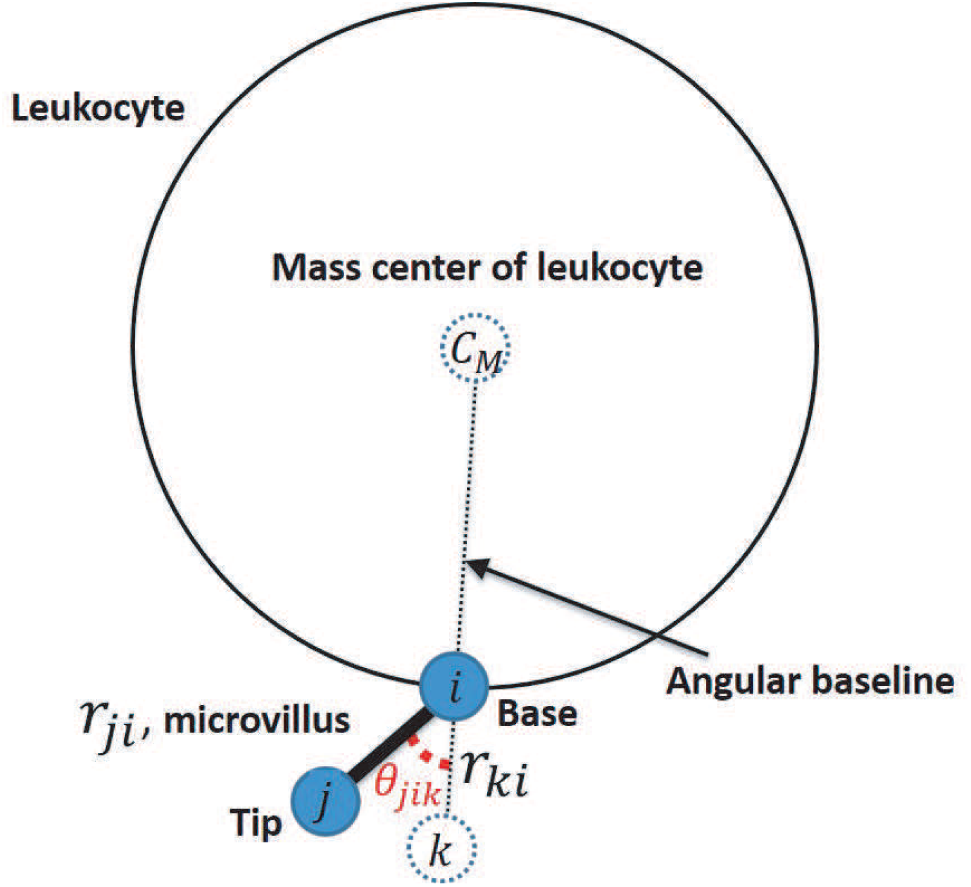
A microvillus is modeled by a spring with its end fixed at the base node *i* on the cell surface. The tip node *j* can be extended or compressed with respect to the base node *i*. In addition, the vector *j* can be bent with respect to vector **r***_ki_* to have a bending angle *θ_kij_*, where the node *k* is always located on the extension portion of the line connecting the mass center *C_M_* to the base node *i*.

Second, vector **r***_ji_* can be bent with respect to vector **r***_ki_* to have a bending angle *θ_kij_*, where *k* is always located on the extension portion of the line connecting the mass center *C_M_* to the base grid *i*. The line is called the angular baseline (the dashed line in Fig. 1) and is used for the measurement of the bending angle only. In fact, **r***_ki_* is a virtual bond, and grid *k* does not participate in simulation. The energy due to the bent bond among three particles 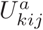 is written by:

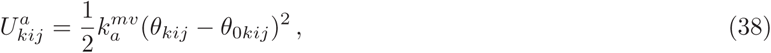

where 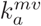 is the angular spring constant, and *θ*_0_*_kij_* represents the equilibrium angle, as shown in Fig. 1. The elastic forces due to the extensional and bending deformation are calculated from the negative gradient of the sum of 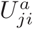 and 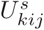 with respect to solid grids *i* and *j*.

Base on previous experimental measurements and numerical studies (9, 24-26), the extensional stiffness and flexural stiffness of the microvilli are 152 − 1340*pN*/*μm* and 4 − *7pN/μm*, respectively. The extensional stiffness *K^E^* is directly equal to the spring constant 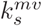, while the flexural stiffness *K* is defined by a ratio of the force on the microvillus to the corresponding deflection, and it is related to the angular spring constant by 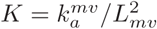 (27). All parameters used in the LSM are given in Table 1 and Table 2.

### Adhesive dynamics

The AD was first presented in a simulation of leukocyte adhesion to endothelial cells by Hammer and Apte (3). This method uses the stochastic Monte Carlo method coupled with kinetics models to simulate the formation and rupture of the receptor-ligand bond.

The kinetics model used in this study is the Dembo model (57), where the forward and reverse rates for the receptor-ligand bond are written as follows:

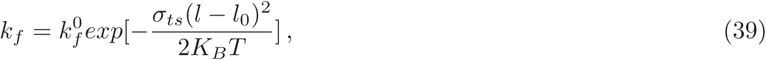

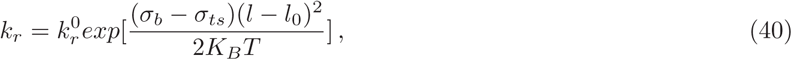

where 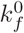 and 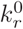 are the unstressed reaction rates; *l* and *l*_0_ re the stretched and unstressed equilibrium bond lengths, respectively; *σ_b_* and *σ_ts_* are the spring constants in the bound and transition states, respectively; *K_B_* is the Boltzmann constant, and *T* is the absolute temperature. The adhesive force *F_b_*, acting on the receptor-ligand bond, is assumed to follow the Hookean spring model, which can be written as:

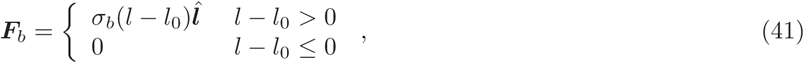

where 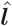 is the unit vector between positions of the receptor-ligand pair.

The probability *P_f_* of formation of a new bond and the probability *P_r_* of rupture of an existing bond in a time interval ∆*t* are given by

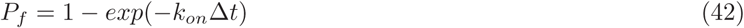

and

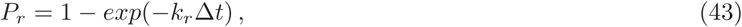

respectively, where *k_on_* = *k_f_ A_L_*(*n_L_ − n_b_*)*; A_L_* is the local surface area on the ligand-coated substrate accessible to each receptor; and (*n_L_ − n_b_*) is the density of the unbound ligand. In the time interval ∆*t*, two random numbers *P_rand_*_1_ and *P_rand_*_2_ between 0 and 1 are generated. A new bond is formed under the condition *P_r_ > P_rand_*_1_. On the other hand, an existing bond is ruptured when *P_r_ > P_rand_*_2_. The time interval ∆*t* of 10*μs* is used, as suggested by Ref. (20), to simulate leukocyte rolling for a period of 3s. All parameters used in the AD are presented in Table 1 and Table 2.

## RESULTS AND DISCUSSION

### Simulation setup

In this study, the leukocyte rolling processes in shear flow are simulated at shear rates of *γ* = 100 − 400*s*^−1^, as shown in Fig. 2. The simulation box is 32 × 46 × 32, where a fluid grid spacing is 5.326 × 10^−7^*m*, and a simulation time step is 1.512 × 10^−7^*s*. A whole leukocyte consists of 714 solid grids for the cell membrane (and also for the microvilli bases), another 714 solid grids for the microvilli tips, and 714 springs for the microvilli. The average membrane particle spacing is 5.326 × 10^−7^*m*, the same as the fluid grid spacing. Two substrates are placed at the bottom and top of the simulation box in the Z-direction. The bottom substrate is at rest and coated with P-selectin at a density of 150/*μm*^−2^. A velocity of *γH_s_* is imposed on the top substrate, where *H_s_* is the distance between the two substrates, and *γ* is the shear rate that varies from 100 to 400*s*^−1^. Periodic boundary conditions are imposed in the X- and Y-directions. Other simulation parameters are given in Table 1. All results shown in this work, except the behaviors of individual microvilli, are calculated by a statistical ensemble average of five repeated independent runs.

**Figure 2:**
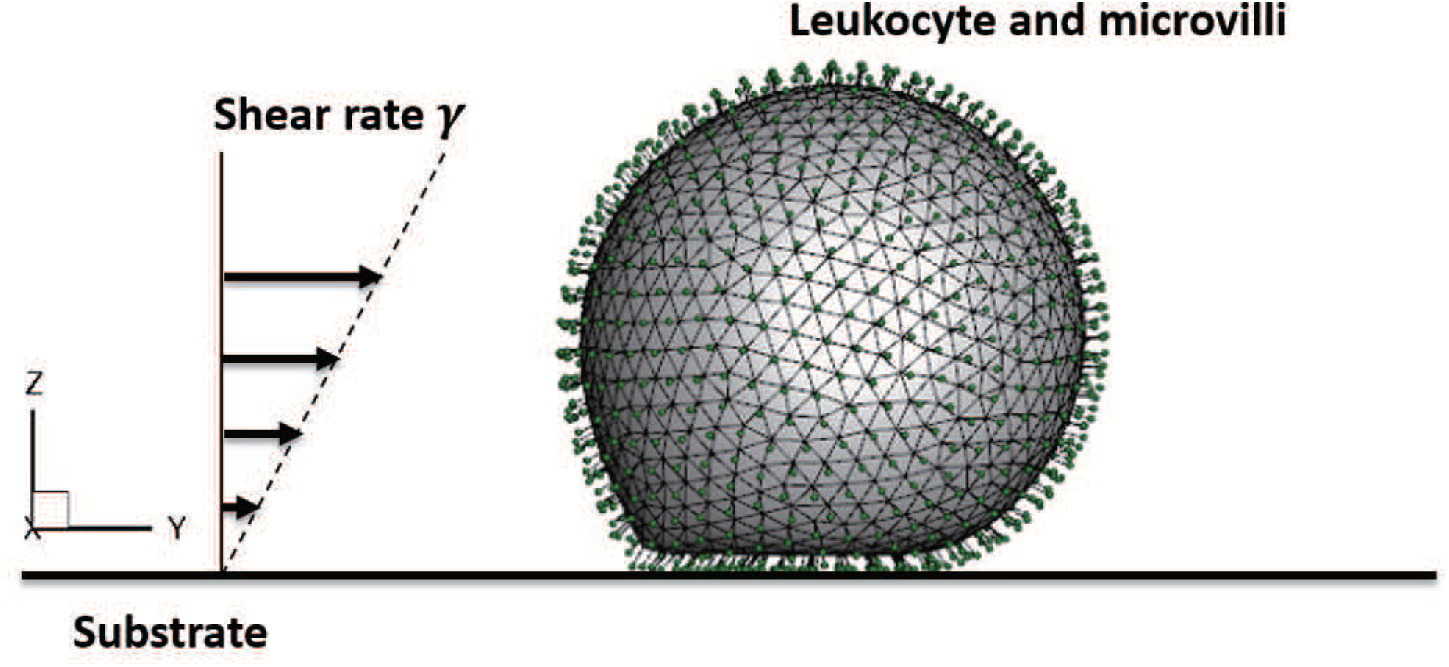
Simulation configuration.

### Validation

For the purpose of validation, a leukocyte rolling on a P-selectin-coated substrate is simulated at a flow shear rate ranging from *γ* = 100*s*^−1^ to 400*s*^−1^. Here, the flexural stiffness of microvilli is set to the experimental value identified by Yao and Shao (24, 25). The simulation results of the average translational velocity of the leukocyte, as a function of shear rate, are compared with those reported by Jadhav et al. (8) and with the experimental data produced by Ramachandran et al. (63) in Fig. 3 (a). Also, the simulation results of the deformation index, defined as leukocyte end-to-end length *L* divided by leukocyte height *H*, and the simulation results of the contact area, between the leukocyte and selectin-coated substrate, are compared with those produced by Jadhav et al. (8) in Fig. 3 (b) and (c), respectively. These results agree with the previously published data in this given range of shear rates. The difference shown by the comparisons can be attributed to distinct numerical models, various simulation constants, and the stochastic nature of the receptor-ligand interaction.

**Figure 3:**
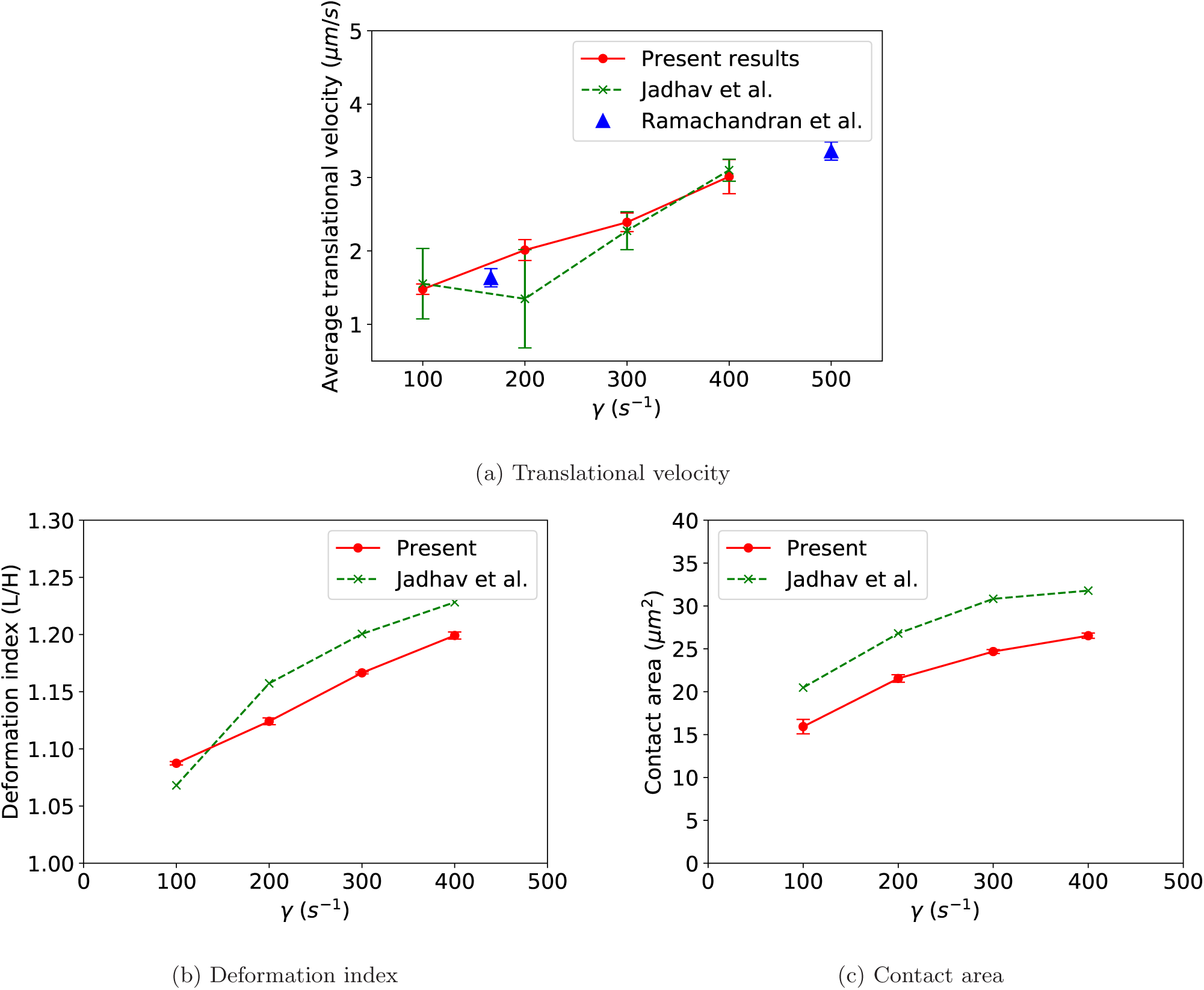
The comparisons between the present simulation results and those of the previously published data from other groups for (a) the translational velocity, (b) the deformation index (*L/H*), and (c) the contact area.

The validation considers not only the rolling velocity, deformation index, and contact area of leukocytes but also concerns their stop-and-go motion, which is one of the distinctive characteristics of leukocyte rollinig. The results of the instantaneous velocity of the leukocyte as a function of time are shown in Fig. 4 (a) for the case of the shear rate of *γ* = 100*s*^−1^ and in Fig. 4 (b) for the case of the shear rate of *γ* = 400*s*^−1^, in which the stop-and-go behavior of the leukocyte rolling is observed. The comparison between Fig. 4 (a) and (b) indicates that shorter time pauses and more frequent peaks with increasing shear rate are consistent with the previous finding by Pappu and Bagch (11).

**Figure 4:**
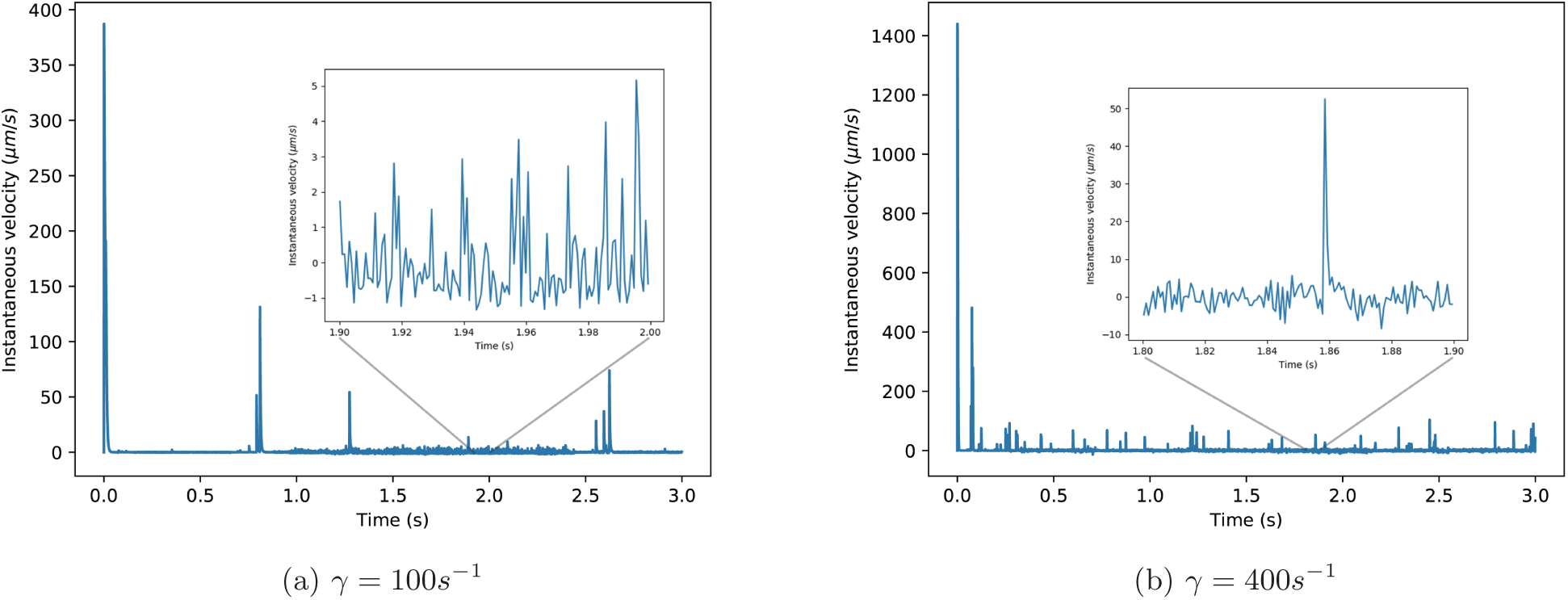
The instantaneous velocity as a function of time at the shear rates of (a) 100*s*^−1^ and (b) 400*s*^−1^. The magnified drawings represent the instantaneous velocity within a 1-second period.

### Effects of the Flexural Stiffness of Microvilli on Rolling Velocity

In order to study the effects of flexural stiffness of the microvilli of a leukocyte or cell on adhesive dynamics, the flexural stiffness varies at three different levels of *K* = 0.1*K_exp_*, *K_exp_*, and 10*K_exp_*, where *K_exp_* = 7*pN/μm* is taken from the experimental data (24, 25), while the extensional stiffness is fixed at *K* = *K^E^* = 1340*pN/μm*, which is adopted from Refs.(7, 14), for all simulations in this work. In this way, the attention is focused on bending deformation only, not on extension, which has been extensively studied by others (17, 19, 21). In this study, the shear rate systematically varies at four different levels: *γ* = 100*s*^−1^, 200*s*^−1^, 300*s*^−1^, and 400*s*^−1^ at each of the given levels of flexural stiffness.

First, the effects of flexural stiffness of microvilli on the leukocyte rolling velocity at different shear rates are probed. The results of an ensemble average of the translational velocity as a function of shear rate at different levels of flexural stiffness is plotted in Fig. 5 (a). It is shown that both the flexural stiffness and shear rate have significant effects on the rolling velocity.

**Figure 5:**
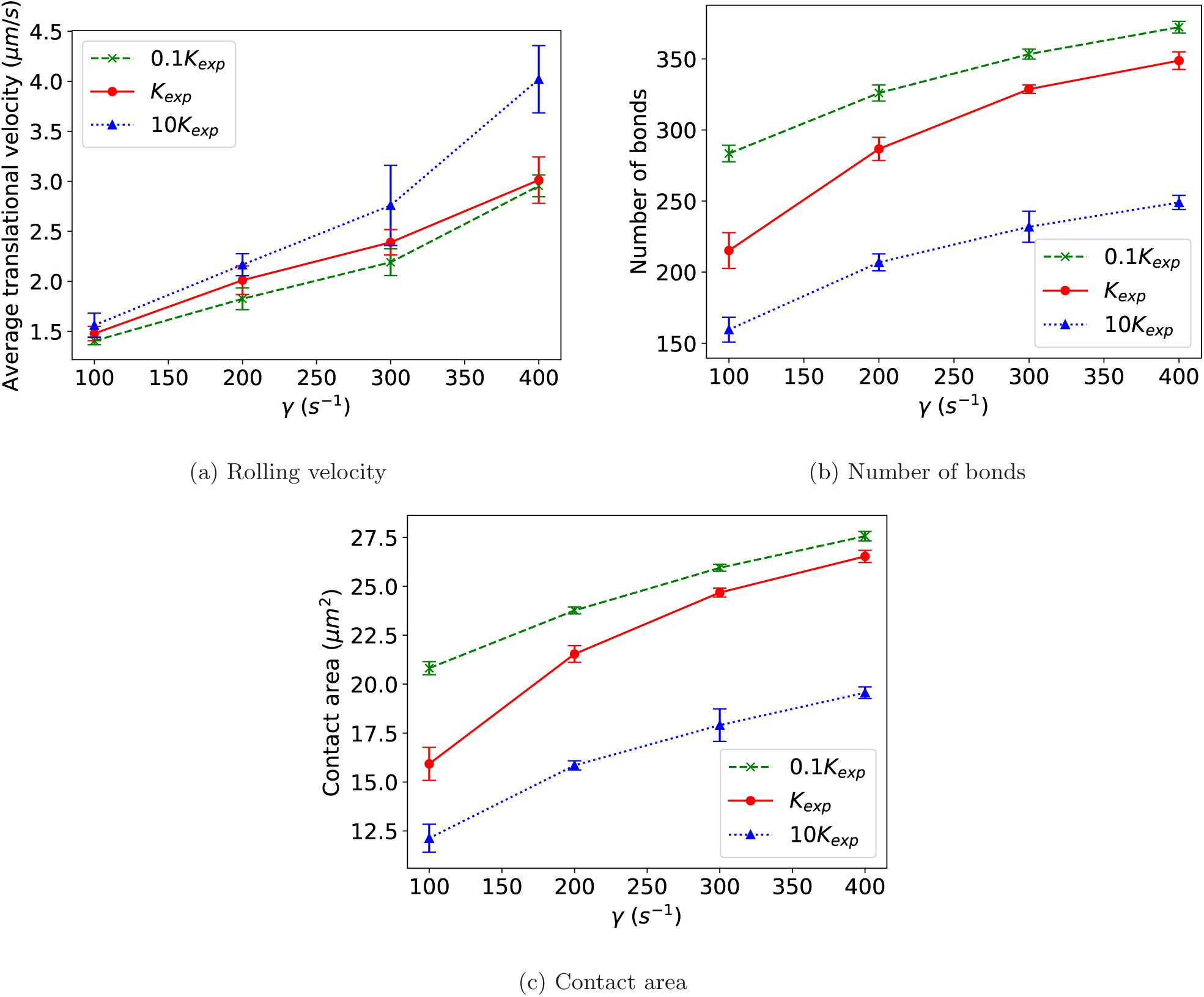
The simulation results of (a) the average translational velocity, (b) number of bonds, and (c) the contact area as a function of a shear rate at three different levels of the flexural stiffness of microvilli. The shear rate is varied from *γ* = 100*s*^−1^ to 400*s*^−1^ at each level of flexural stiffness.

At each of the given levels of flexural stiffness, the rolling velocity increases as the shear rate increases since a larger shear force drives cells to move faster and simultaneously induces a larger number of bonds (see Fig. 5 (b)) and an expanded contact area (see Fig. 5 (c)). The bonds and contact area dynamically respond to the external shear force. The results indicate that the slopes of the rolling velocity curves are larger in the large shear rate range than in the small shear rate range, revealing that the rolling velocity increases faster in the large shear rate range than in the small shear rate range. On the contrary, the slopes of curves of the number of bonds and the contact areas are smaller in the large shear rate range than in the small shear rate range. As expected, the contact areas and the number of bonds are developed more slowly at the large shear rate range, so that the rolling velocity increases faster.

It is also observed that the degree of the increase in the rolling velocity due to the flexural stiffness depends on the shear rate. The rolling velocity increases from 2.95*μm/s* to 4.02*μm/s* at the shear rate of *γ* = 400*s*^−1^, while the corresponding velocity increases from 1.41*μm/s* to 1.56*μm/s* at the shear rate of *γ* = 100*s*^−1^. The net increase in rolling velocity due to the flexural stiffness is approximately 7 times larger at *γ* = 400*s*^−1^ than at *γ* = 100*s*^−1^, although they have the same levels of flexural stiffness.

At a given shear rate, the rolling velocity increases as the flexural stiffness of the microvilli increases or as the flexibility of microvilli decreases, because the microvilli with a larger flexural stiffness have a smaller bending deformation and are likely to be oriented more vertically on the cell surface. Thus, the contact area and number of bonds between the cell and the substrate all become smaller, making the cell roll faster, as compared to cells with more flexible microvilli. The results of the number of bonds and the contact areas are viewed in Fig. 5 (b) and (c), where the number of bonds and the contact areas dramatically decrease as the flexural stiffness increases.

Next, the results of the deformation index as a function of the shear rate are given in Fig. 6. The figure shows that at each of the given levels of flexural stiffness, the deformation index increases as the shear rate increases, an outcome which is consistent with the previous findings by Jadhav et al. (8). The deformation index increases by 10 − 11% when the shear rate increases from *γ* = 100*s*^−1^ to *γ* = 400*s*^−1^. However, at each of the given shear rates, when the flexural stiffness is largely varied from *K* = 0.1*K_exp_* to *K* = 10*K_exp_*, the deformation index has only 1 – 3% variation, illustrating that the cell bulk deformation is more sensitive to the shear rate than the flexural stiffness. In other words, the bulk cell deformation strongly depends on the shear rate and less significantly on the flexural stiffness.

**Figure 6:**
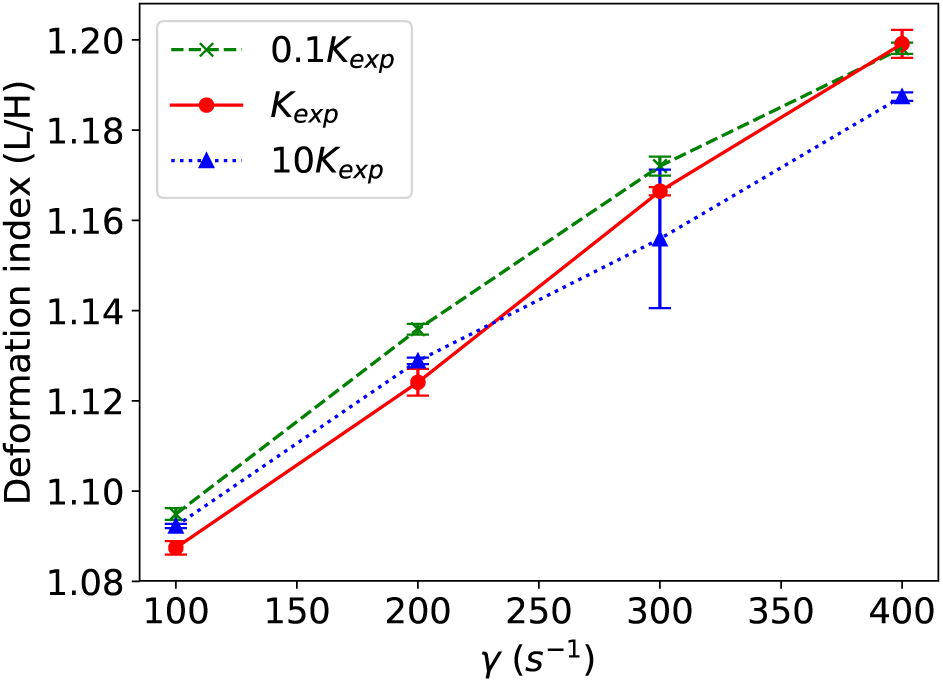
The simulation results of the deformation index at three different levels of the flexural stiffness of microvilli at the shear rates from 100*s*^−1^ to 400*s*^−1^.

For the purpose of visualization, the animations of leukocyte rolling at shear rates of *γ* = 100*s*^−1^ and 300*s*^−1^ are presented in movie S1 and movie S2 in the Supporting Material.

Subsequently, Fig. 7 illustrates the instantaneous velocity of leukocyte as a function of time at three different levels of the flexural stiffness of microvilli at the shear rate of *γ* = 300*s*^−1^. As the flexural stiffness increases, the time pauses become shorter, peaks are higher, and the rolling velocity in the case of *K* = 0.1*K_exp_* experiences more frequent fluctuations between peaks, due to more occurrences of bonding and de-bonding, as compared to the velocities in the cases of *K* = *K_exp_* and *K* = 10*K_exp_*.

**Figure 7:**
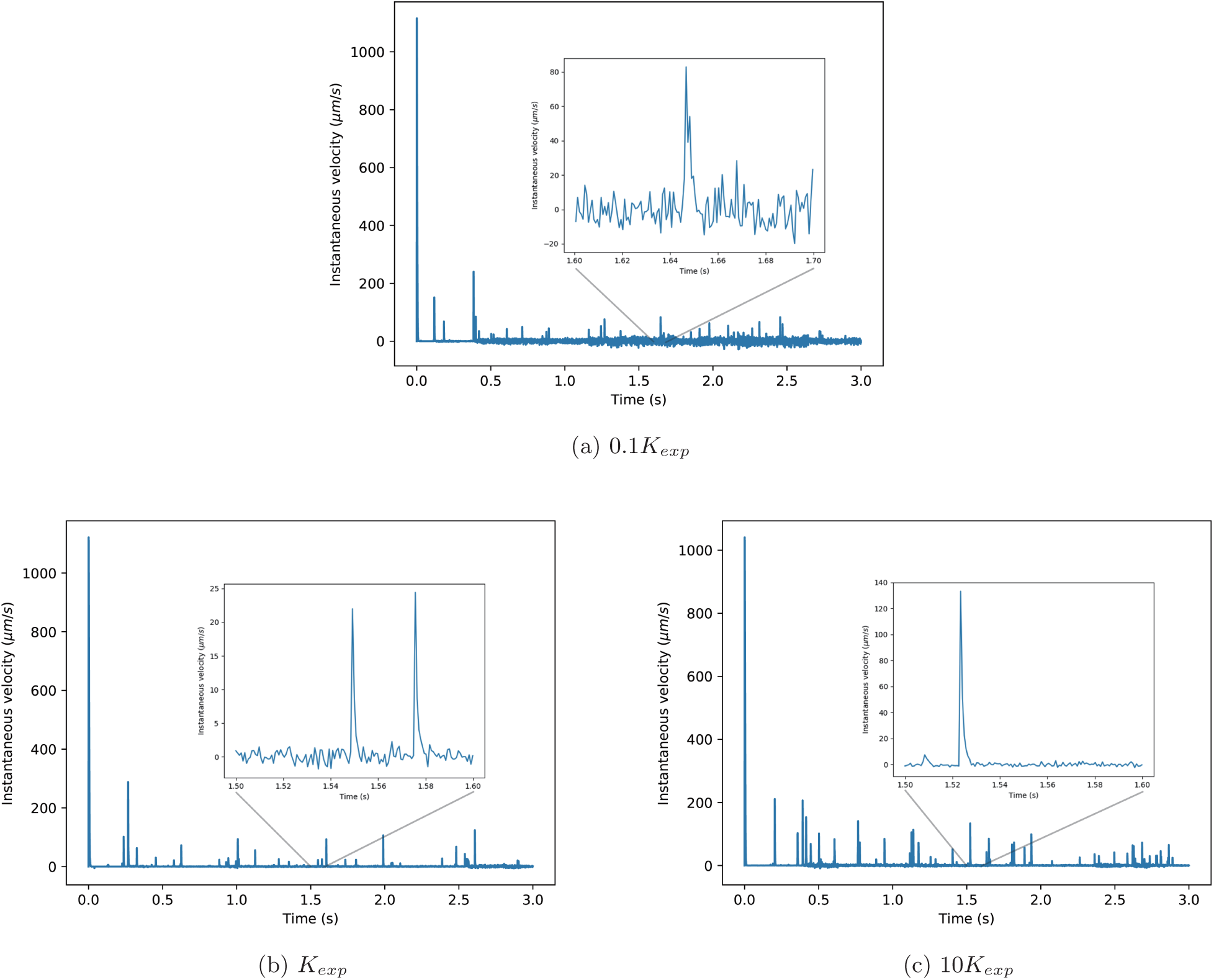
TThe instantaneous velocity at microvillus flexural stiffness of (a)0.1*K_exp_*, (b)*K_exp_*, and (c)10*K_exp_* at the shear rate of *γ* = 300*s*^−^1. The magnified drawing presents the instantaneous velocity in 0.1-second period.

### Effects of the Flexural Stiffness of Microvilli on Bond Forces

A total adhesive bond force is based on the global coordinates of X, Y, and Z and defined by

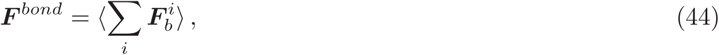

where *i* denotes the *i*th bond.

In order to distinguish bending force from extensional force, a bond force can be decomposed into two components in a local coordinate system associated with a microvillus: one is parallel to the direction of the angular baseline, and the other is perpendicular to it. The parallel component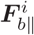 is responsible for the extensional deformation, and the perpendicular component 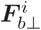is responsible for the bending deformation, where *i* denotes the *i*th bonded microvillus.

To consider the collective effects of the bond forces on all microvilli and to separate the role of the bending force from that of extensional force, total parallel and perpendicular force components based on the local coordinates are defined by

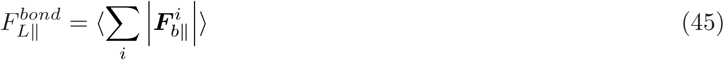

and

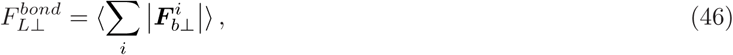

respectively. The corresponding total local coordinate-based bond force 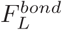 is defined by

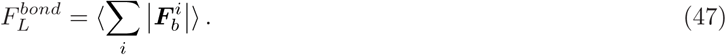

The total adhesive bond force ***F****^bond^*, total local coordinate-based bond force 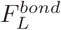, its parallel component 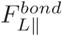, and perpendicular component 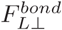 are computed at three different levels of flexural stiffness, and the results as a function of shear rate are displayed in Fig. 8 (a), (b), (c), and (d), respectively. At each of the given levels of the flexural stiffness, the results of all the bond forces increase monotonically as the shear rate increases. This occurs because the adhesive bond force should be balanced by the hydrodynamic shear force, which drives the cell to move and induces a responding adhesive bond force. The larger the shear rate, the larger the responding adhesive bond force. Significantly, the bond force strongly depends on shear rates. In this study, the Reynolds number is *R_e_* ∈ [1.172 × 10^−3^, 4.688 × 10^−3^], and therefore the inertial effect can be neglected.

**Figure 8:**
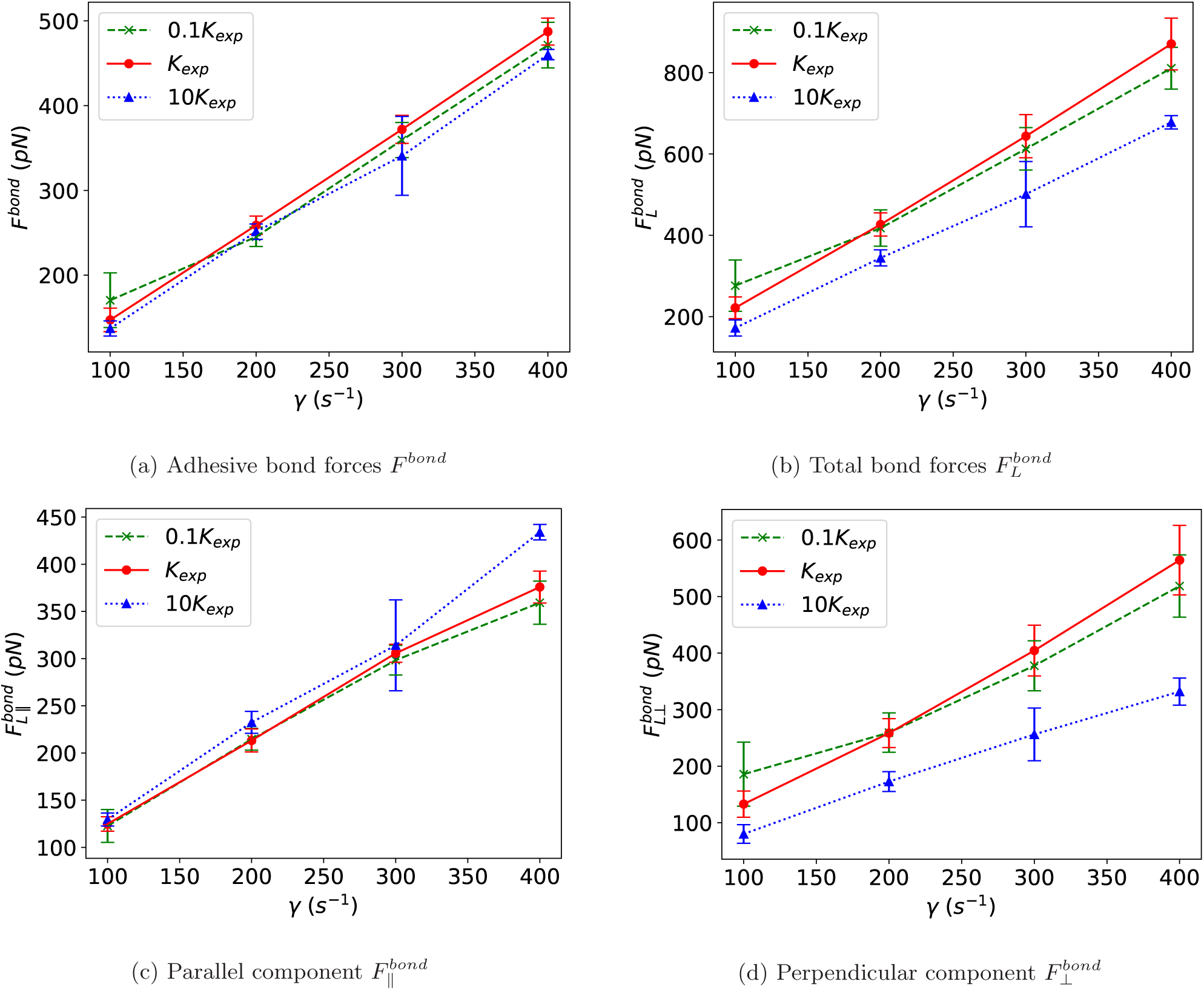
The simulation results of (a) the adhesive bond force, (b) the local-coordinate-based total bond forces 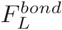, (c) the local-coordinate-cased parallel component 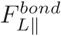, and (d) the local-coordinate-based perpendicular component 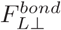, at three different levels of the flexural stiffness at the shear rates from *γ* = 100 to 400*s*^−1^.

The study confirms that the bond force is influenced not only by the shear rate but also by the flexural stiffness. At the given lowest shear rate of *γ* = 100*s*^−1^, the result of the bond force increases monotonically as the flexural stiffness decreases and arrives at a maximum at the lowest level of the flexural stiffness of *K* = 0.1*K_exp_*. This result is due to the largest bending deformation of microvilli (see the next section), which may result in a larger contact area and a larger bonding probability. At the given highest level of the shear rate of *γ* = 400*s*^−1^, the bond force increases first as the flexural stiffness decreases from *K* = 10*K_exp_* to *K_exp_* and then decreases as the flexural stiffness continuously decreases to the level of *K* = 0.1*K_exp_*. The maximum bond force occurs at an intermediate level of the flexural stiffness. A similar behavior is observed at *γ* = 200*s*^−1^ and 300*s*^−1^. It is observed that across the entire shear rate range, the amplitude of the increase in ***F****^bond^* and 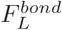 is largest for the case of the intermediate flexural stiffness of *K* = *K_exp_*. If the flexural stiffness is very small (*K* = 0.1*K_exp_*) or the microvilli are too flexible, the bending deformation is saturated even at the small shear rate and lends to a negligible difference when compared to the large shear rates. Similarly, if the flexural stiffness is very large or the microvilli are too stiff, the bending deformation lends to a negligible difference across all the shear rates. Importantly, the maximum amplitude of the increase of the bond force occurs at the flexural stiffness of the experimental value of *K* = *K_exp_*. It is inferred that an interplay of the roles between the flexural stiffness and flow shear rate determines the final total bond force.

As shown in Fig. 8 (d), the perpendicular component forces are profoundly affected by the flexural stiffness at each of the given shear rates. For example, at the lowest level of the shear rate of *γ* = 100*s*^−1^, the perpendicular component increases from 80*pN* to 185*pN*, as the flexural stiffness continuously decreases from *K* = 10*K_exp_* to *K* = 0.1*K_exp_*, while the parallel component just slightly changes from 122*pN* to 129*pN*, as shown in Fig. 8 (c). The net force change due to the flexural stiffness is 15 times larger in the perpendicular component than in the parallel component. It is concluded that the change in the total bond force primarily results from the change in the perpendicular component. This outcome can also be seen from the case of the shear rate of 400*s*^−1^. The perpendicular component increases from 331*pN* to 518*pN*, while the parallel component changes only from 433*pN* to 359*pN*. Again, the increase in the perpendicular component is much larger than the change in the parallel component due to the bending of microvilli. In fact, the bending angle significantly increases as the flexural stiffness decreases or as the flexibility increases, as discussed in the next section.

### Bending Deformation of Microvilli

In order to examine the influence of the flexural stiffness of microvilli on their bending deformation, an angular distribution function *f*(*θ*) of the bounded microvilli is defined by

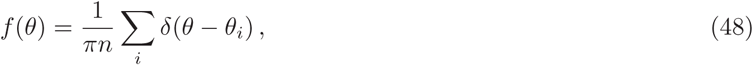

where *n* is the total number of the bonded microvilli, and *θ_i_* is the bending angle of the *i*th bonded microvillus. The angular distribution function is utilized to describe the probability of finding a microvillus at an angle of *θ* per unit angle.

The results of an ensemble average of the angular distribution function 〈*f*(*θ*)〉, as a function of the bending angle at three different levels of the flexural stiffness, are displayed and compared in Fig. 9, at each of the four different shear rates, where the green dashed, red solid, and blue dash-dot lines represent the data sets corresponding to *K* = 0.1*K_exp_*, *K_exp_*, and 10*K_exp_*. It is shown that the flexural stiffness has a great impact on the bending deformation of microvilli. For example, for the case of the shear rate of *γ* = 100*s*^−1^, three curves of the angular distribution function are compared in Fig. 9 (a). One sharp peak appears on the curve of the largest flexural stiffness of *K* = 10*K_exp_*. This peak value is as high as 0.95 and is located near the angle of zero, evidencing that most microvilli are oriented in the direction perpendicular to the cell surface or along the angular baseline. As the flexural stiffness reduces to a lower level of *K* = *K_exp_*, two peaks are observed: one peak with a value of 0.18 is located near *θ* = 0, and the other with a similar value of 0.18 is located at *θ* = 2*π*/9. As the flexural stiffness continuously reduces to the lowest level of 0.1*K_exp_*, one wider and lower peak appears and shifts to the right side closer to the angle of *θ* = *π*/2, indicating that as the flexural stiffness reduces, the bending angle increases, and the microvilli tend to be flattened on the cell surface. A similar behavior is observed at every given shear rate. It is pointed out that for a given flexural stiffness, the peak becomes lower and wider and shifts to the side of the angle of *θ* = π/2 as the shear rate increases, suggesting that more microvilli are moved from the angular baseline direction to their perpendicular direction.

**Figure 9:**
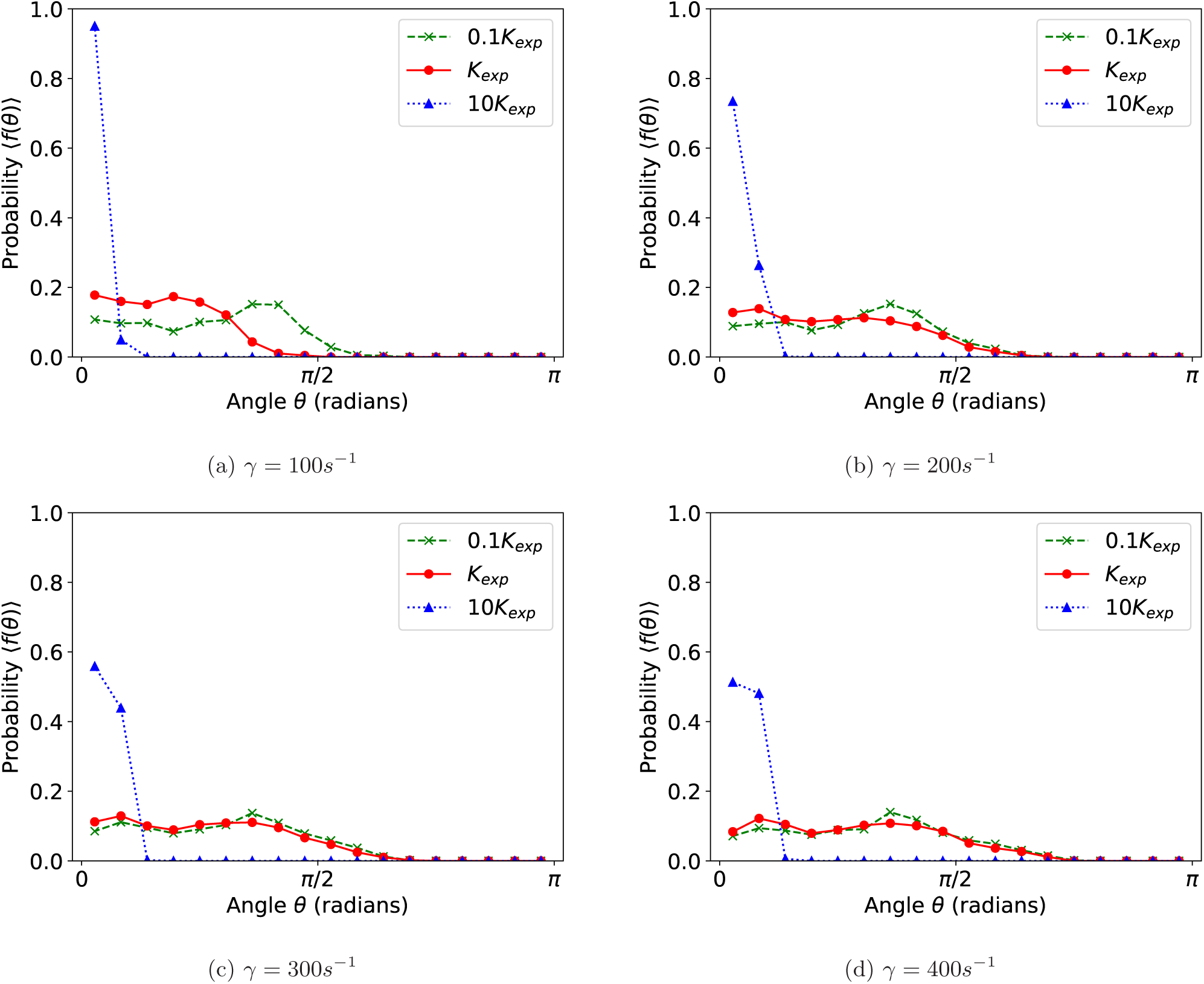
The simulation results of the angular distribution of the microvilli at three different levels of the microvillus flexural stiffness at shear rates of (a) 100*s*^−1^, (b) 200*s*^−1^, (c) 300*s*^−1^, and (d) 400*s*^−1^.

### Bending Behavior of Individual Microvillus

To closely monitor the process of bonding formation at the microscopic scale of a single microvillus during cell rolling, four individual microvilli are selected, and their bending angles as a function of time are recorded as a sample. Fig. 10 depicts the angles of the four individual microvilli as a function of time, where the yellow solid lines and blue dotted lines represent the duration involved in bonded and non-bonded states, respectively. Each figure also includes both the side and bottom views of the leukocyte. For the bottom view, the red marker specifies the particularly selected single microvillus (e.g., microvilli No. 1, 2, 3, and 4), and the black points denote the other microvilli. Microvillus No. 1 is initially located near the center of the contact area, while No. 2 is initially located near the right edge of the contact area. The same shear rate *γ* = 400*s*^−1^ is applied on microvilli No. 1 and 2. Microvilli No. 3 and 4 are similar to No. 1 and 2, except that the smaller shear rate *γ* = 100*s*^−1^ is used.

**Figure 10:**
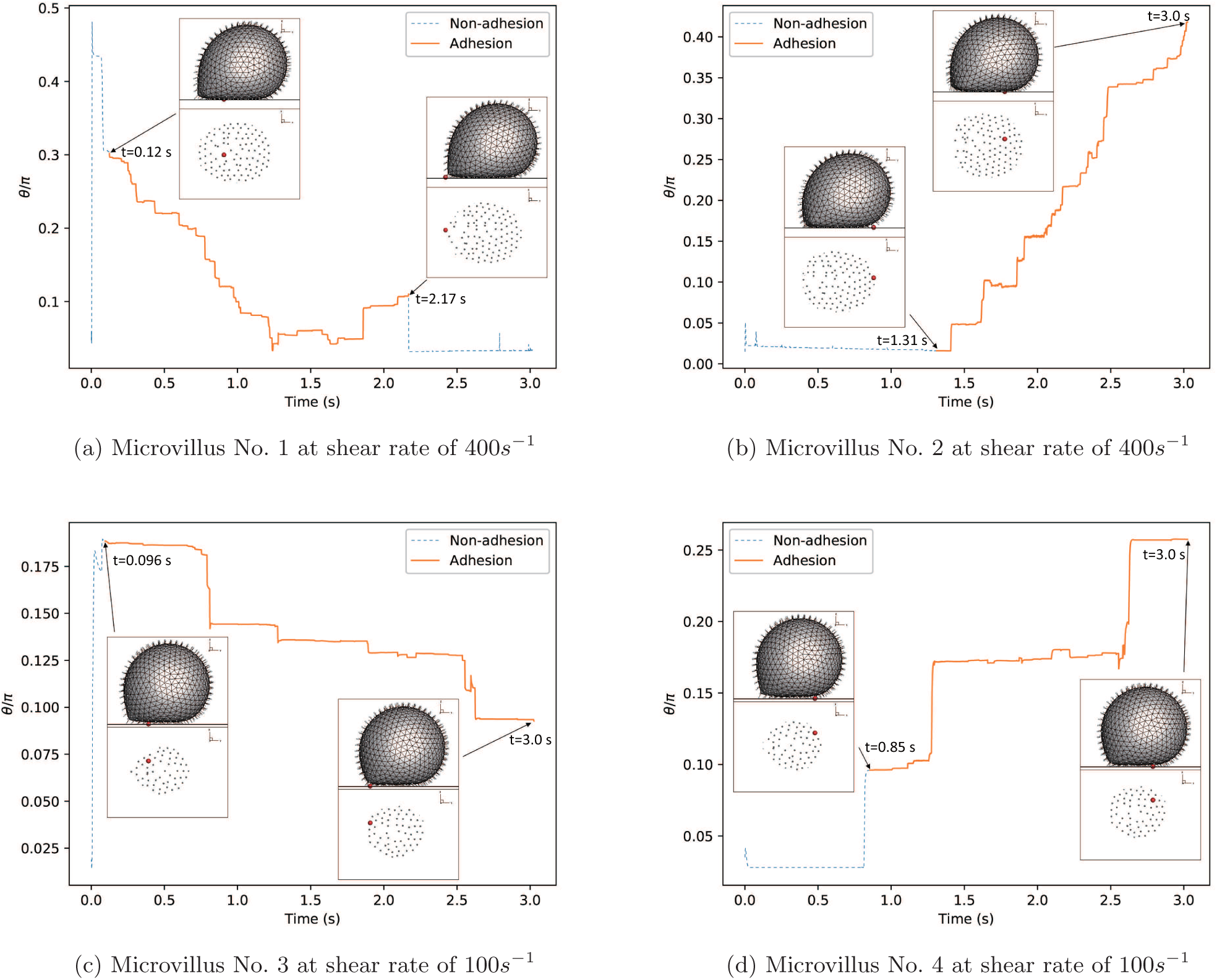
The bending angle as a function of time for (a) the single microvillus initially located near the center of the contact area at *γ* = 400*s*^−1^, (b) the single microvillus initially located near the right edge of the contact area at *γ* = 400*s*^−1^, (c) the single microvillus initially located near the center of the contact area at *γ* = 100*s*^−1^, and (d) the single microvillus initially located near the right edge of the contact area at *γ* = 100*s*^−1^. The stiffness of the microvillus is fixed at the experimental value.

The variation of the bending angle of microvillus No. 1 is shown in Fig. 10 (a) as the microvillus moves from the center to the left edge of the contact area during the rolling process. In the initial period (*t <* 0.12s), there is non-bonding involved (blue dotted line). The microvillus stands vertically on the cell surface with a very small bending angle and then is quickly bent with a large angle and flattened with the surface. Soon after that, a receptor-ligand bond is developed and formed at *t* = 0.12s. As the leukocyte continues to roll, the angle of the microvillus first decreases and then increases. Finally, the bond is broken at *t* = 2.17s and the microvillus becomes vertical again with a small bending angle that occurs when the microvillus moves to the left edge of the contact area. This non-bonding, bonding, and de-bonding process corresponds to the stop-and-go motion. The bending behavior of microvillus No. 2, moving from the right edge to the center of the contact area during the rolling process, is shown in Fig. 10 (b). A bond is formed on the vertical microvillus at *t* = 1.31s. Unlike No. 1, the bending angle increases monotonically, and this microvillus becomes more flattened with the cell surface when it migrates to the center of the contact area.

Over all, the behaviors of microvilli No. 3 and 4 in Fig. 10 (c) and (d) are similar to No. 1 and 2. The major difference is that the bending angles of No. 3 and 4 are smaller, as expected, due to the smaller shear rate *γ* = 100*s*^−1^. The rupture of bonding is observed in micrivillus No. 1 only.

## CONCLUSIONS

The details of the LLM, including LBM, IBM, CGCM and AD, are described in this manuscript for the first time. This developed method can treat not only extensional but also bending deformation of microvilli of a leukocyte or cell in flows. The effects of flexural stiffness of microvilli on rolling adhesive process in a low-Reynolds-number shear flow at different shear rates are simulated and studied by using the LLM. The flexural stiffness is varied at three different levels, while the extensional stiffness is fixed at a constant, allowing one’s attention to be focused on the effects of flexural stiffness only.

At each given level of the flexural stiffness, the flow shear rate is varied at four different levels, and the results of the bond force, number of bonds, contact area, bending angular distribution, and cell deformation index are computed. To address the bending, the bond force is decomposed into two directions, parallel and perpendicular to the direction of cell surface in an associated local coordinate, so that the perpendicular force component is separated from the parallel force component, and the role of the perpendicular force component on bending is extracted and analyzed.

Several interesting behaviors are revealed from simulations:

1. At each of the given levels of the flexural stiffness, the rolling velocity, total bond force, number of bonds, bending angles, and contact area increase as the shear rate increases.
2. At a given shear rate, the flexural stiffness also significantly affects the bond force, bending deformation, number of bonds, and contact area. The angular distribution function is employed to statistically measure the bending deformation. The position of the peak of the curve of the angular distribution function of the microvilli shifts to the side of *θ* = *π*/2, the value of the peak reduces, and the peak becomes wider and lower, as the stiffness reduces, illustrating that the microvilli become more flattened with the surface and have a larger probability to be bonded with the substrates.
3. The interplay of the roles between the shear rate and flexural stiffness of microvilli determines the final bond force.
4. It is demonstrated that the bending deformation and perpendicular force component strongly depend on the flexural stiffness, while the bulk cell deformation mainly depends on shear rates and is not sensitive to the flexural stiffness of microvilli.

The above information may improve the efficacy of diagnoses and treatments, especially within the realm of technological advances, such as the design of microfluidic chips.

## SUPPLEMENTARY MATERIAL

Two movies are available at http://www.biophysj.org.

## ACKNOWLEDGMENTS

Wu and Qi thank Dr. M. Di Pierro and Yi-Chin Huang for proofreading and writing advise. Wu also acknowledges the Gwen Frostic Doctoral Fellowship for partially supporting this work and Qi acknowledges the award of Faculty Creativity Initiative of College of Engineering and Applied Sciences at the Western Michigan University.

